# Single-Allele Chromatin Tracing Reveals Genomic Clustering of Paralogous Transcription Factors as a Mechanism for Developmental Robustness in T Cells

**DOI:** 10.1101/2025.05.30.656885

**Authors:** Atishay Jay, Yeqiao Zhou, Sora Yoon, Bereketab N. Abeje, Aditi Chandra, Julia Wald, Arjun Raj, Robert B. Faryabi, Golnaz Vahedi

## Abstract

In metazoans, gene duplication has given rise to paralogous transcription factors, which have functionally diversified to control cellular differentiation. While the majority of paralogous TFs are dispersed across different chromosomes, some remain clustered raising the question of whether genomic proximity confers any evolutionary advantage for TF clusters. To address this, we investigated a ∼1 Mbp locus containing two ETS family paralogs, *Ets1* and *Fli1*. Using a sub-diffraction sequential imaging technique called Optical Reconstruction of Chromatin Architecture (ORCA), we traced the 3D organization of this region in single alleles of T cells from genetically engineered mice with targeted deletions of key regulatory elements. In wild-type T cells, the predominant chromatin conformation spatially links *Ets1* to its proximal super-enhancer, segregating *Ets1* from *Fli1*. This topology correlates with high *Ets1* and low *Fli1* expression. Deletion of the *Ets1* super-enhancer abolishes this configuration, triggering locus-wide architectural rewiring that increases *Ets1-Fli1* promoter-promoter interactions and subsequently the co-expression of two genes within individual cells. Remarkably, this compensatory interaction bypasses insulated chromatin domains, sustaining *Ets1* levels necessary for T cell development despite enhancer loss. Our results reveal that genomic clustering of TF paralogs enables dynamic architectural plasticity: while a super-enhancer fine-tunes paralog expression balance in wild-type contexts, its deletion unmasks latent promoter-driven coordination, suggesting that proximity safeguards functional redundancy and transcriptional resilience critical for cellular fitness.

## Main

Transcription factors (TFs) orchestrate complex cellular processes by interpreting genomic information encoded in DNA^1^. Gene duplication events, including two rounds of whole-genome duplication at the base of vertebrate evolution in addition to tandem and segmental duplications has largely expanded TF diversity^2–4^. These duplication events have given rise to paralogous TFs, which have acquired specialized functions, fine-tuning gene regulation and cell fate determination^5^. In the mammalian genome, most paralogous TFs from the same family are located on different chromosomes rather than being physically clustered. However, notable exceptions exist in which paralogous TFs are positioned in close proximity on the same chromosome^6–9^. This raises a question: does the linear genomic arrangement of certain paralogous TFs confer any evolutionary advantage? In *Drosophila*^10, 11^ and zebrafish^12^, studies have proposed a model in which genomic proximity of paralogous genes enhances coordinated transcription by enabling sharing of regulatory elements. Whether this paradigm is generalizable in the mammalian genomes or whether the close genomic arrangement of paralogous TFs serves alternative functional or evolutionary roles remains unclear.

The E26 transformation-specific (ETS) family of TFs represents an ancient and evolutionarily conserved group of paralogous TFs, with their ancestor tracing back to earliest metazoans^13^. There are approximately 28 distinct ETS paralogs identified in the mammalian genome^14, 15^. Notably, four paralogous pairs, *ETS1* and *FLI1*, *ETS2* and *ERG*, *ELF5* and *EHF*, in addition to *ETV3* and *ETV3L,* are located in genomic proximity albeit on different chromosomes. Yet, the functional importance of ETS TF pairing in regulating gene expression and cellular processes is unclear. In this study, we investigated the three-dimensional (3D) genome organization of a ∼1Mbp genomic locus harboring the *Ets1-Fli1* TF pair along with multiple enhancers and CTCF-bound domain boundaries. We chose to study this locus in T cells based on several key features: (1) the formation of a “multi-enhancer hub”^16–18^, demonstrating extensive enhancer-enhancer interactions in T cells according to population-level genomic assays^19^; (2) the unusual association of this locus to T cell mediated diseases including allergy, asthma, lupus, and rheumatoid arthritis^20, 21^; (3) the stretches of highly acetylated histones defined as super-enhancers, flanked by two repressive H3K27me3 domains; and (4) the distinct expression profiles and functional significance of *Ets1* and *Fli1* in T cells^22, 23^.

Population-level assays that fragment the chromatin and capture only pairwise interactions fail to provide a complete view of 3D chromatin architecture. To unravel the *cis* regulation of *Ets1* and *Fli1* TF pair positioned within a multi-enhancer hub, we need to employ a technology capable of tracing the chromatin fiber in individual cells, capturing the full spectrum of chromatin folding configurations for both genes and their regulatory elements at the same time in the same allele.

Here, we implemented a sequential imaging approach with sub-diffraction resolution called Optical Reconstruction of Chromatin Architecture (ORCA)^24^, which enabled tracing of the *Ets1-Fli1* locus at the single-allele level. In order to study the significance of genomic proximity between the *Ets1-Fli1* paralogous pair, we generated two genetically engineered mouse strains in which DNA sequences corresponding to a super-enhancer proximal to the *Ets1* gene^20^ and a CTCF boundary element were separately deleted. We performed single molecule RNA FISH in addition to chromatin tracing experiments in mature T cells of the thymus from these mouse strains. In wild-type T cells, the most prevalent chromatin conformation brings *Ets1* into spatial proximity with its super-enhancer, segregating *Ets1* from its paralog *Fli1*. This frequent genome topology corresponds to cells expressing high levels of *Ets1* and low levels of *Fli1* transcripts (*Ets1^high^Fli1^low^*). Deleting the super-enhancer proximal to *Ets1* leads to the disappearance of *Ets1^high^Fli1^low^* cells. The entire locus rewires after the super-enhancer deletion, leading to an increase in *Ets1*-*Fli1* promoter-promoter interactions and co-expression of the two paralogs within individual cells. This topology occurs despite *Ets1* and *Fli1* positioning in two insulated domains, suggesting border bypassing^25^ occurs frequently when the *Ets1* super-enhancer is not pulling *Ets1* from *Fli1*. The consequence of border bypassing after genetic loss of *Ets1* super-enhancer is maintaining *Ets1* levels within a tolerable range required for T cell development. We also deleted a boundary element bound by CTCF downstream of *Ets1* which led to the expansion of the entire locus. Together, these findings demonstrate that the *Ets1* super-enhancer orchestrates allele-specific chromatin architecture to balance paralog expression, and its deletion unveils a compensatory promoter-promoter interaction network essential for maintaining *Ets1* levels critical to T cell development.

## Results

### Most paralogous TFs are genomically isolated from other family members

Eukaryotic genomes exhibited duplication events in TF genes, leading to paralogous TFs sharing highly similar DNA-binding domains that characterize specific TF families. This prompted us to ask: how frequently paralogous TFs are genomically isolated from other members within their respective TF families compared with being clustered together on the same chromosome? We defined “clustered paralogous TFs” as TFs from the same family located within a 300 kb window on the same chromosome. Among the 1,637 TFs in the human genome^26^, 1,058 TFs are genomically isolated from other members of their families (Figure 1a). These include key regulators of cell identity and function, such as *TCF7, EBF1, FOXO1, ID2,* and *MYC*. The remaining 579 TFs are found in genomic clusters. Notably, 72% (422) of these clustered TFs belong to a single TF family—the C2H2 zinc finger (ZNFs) proteins. ZNFs with KRAB domains bind and silence transposable elements and comprise the largest family of mammalian TFs, rapidly evolving within and between species^27^. In fact, the largest TF cluster in the human genome spans 1.4 Mbp on chromosome 19 and contains over 50 KRAB-ZNF genes (Figure 1b). Additional large clusters include members of the *HOXA* (12 genes) and *HOXD* (10 genes) families. Beyond these exceptions, the most frequent genomic clustering of paralogous TFs in the human genome occurs as isolated pairs where two paralogous TFs are located in genomic proximity on the same chromosome, likely originating from ancient tandem duplication events (Figure 1a-b, Table S1).

**Figure 1.**
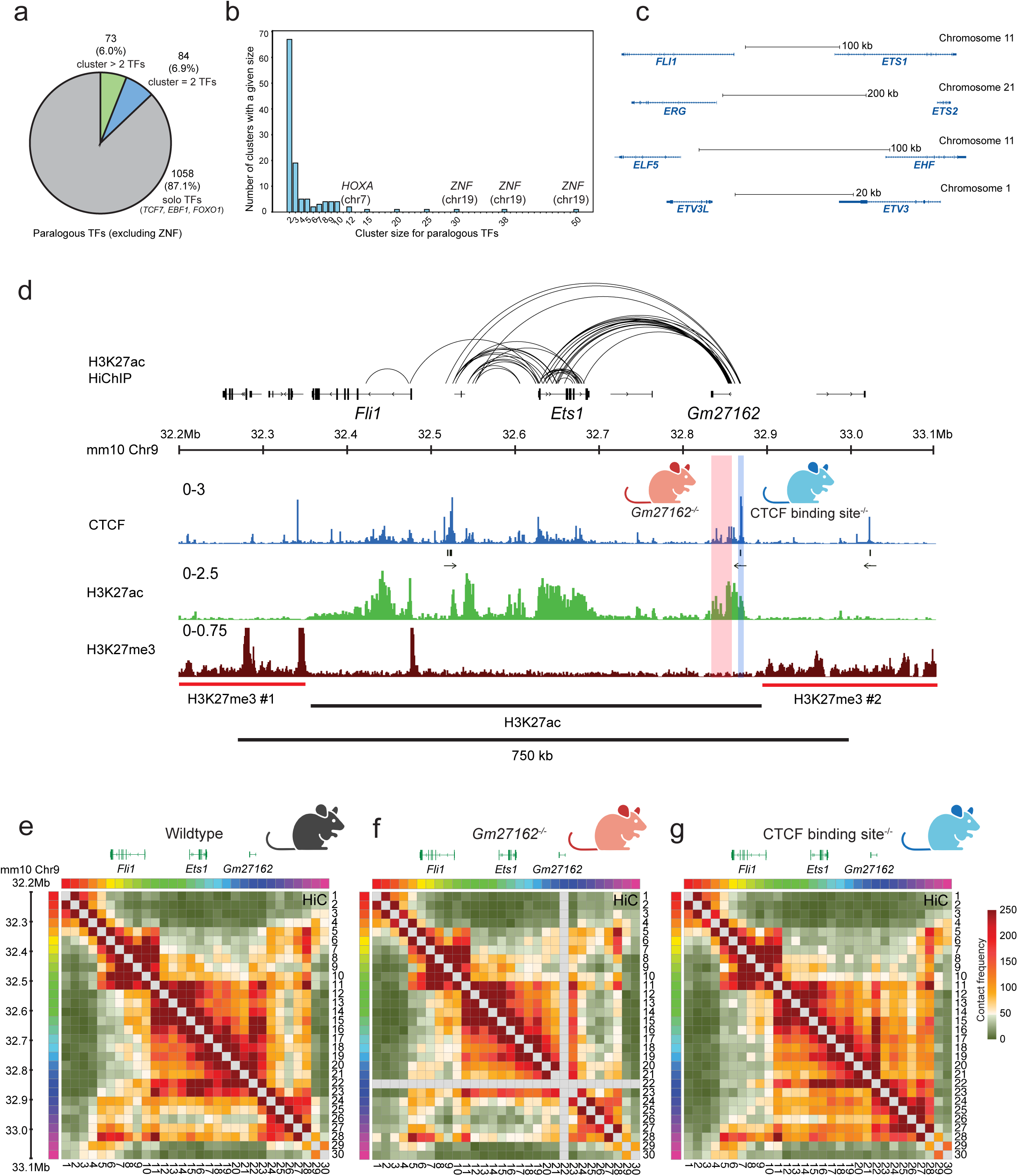
Enhancer connectivity at the *Ets1-Fli1* locus is flanked by H3K27me3-rich repressed chromatin. **(a)** Proportion of paralogous transcription factor (TF) genes clustered based on physical proximity threshold of 300 kbp. Grey region of the pie chart shows ‘Solo TFs’ which denotes TFs in the genome without any paralogous TF in 300kbp on the same chromosome (e.g., *TCF7, EBF1, FOXO1*). The blue region represents clusters of paired paralogous TFs and the green region represents TF clusters with more than two TFs with the exclusion of the C2H2 zinc finger (ZNFs) proteins TFs. **(b)** Distribution of clustered paralogous TF genes based on cluster size. The x-axis represents the cluster size where paralogous TF genes are defined to cluster if they are within 300 kbp of each other on the same chromosome. The y-axis represents the frequency of observing a specific cluster size. High cluster size TFs such as *ZNF* and *HOXA* are labeled. **(c)** Four E26 transformation-specific (ETS) family paralogous TF pairs annotated by chromosome position and proximity between TF genes in the human genome. **(d)** Genome browser views show H3K27ac and H3K27me3 modifications at the *Ets1-Fli1* locus measured by CUT&RUN. The 750 kbp linear genomic distance between the H3K27me3 domains is denoted by the horizontal black bar. CTCF CUT&RUN and motif analysis show the magnitude and directionality of CTCF binding. High degree of interactions within the *Ets1-Fli1* locus is shown by H3K27ac HiChIP. The 25kbp deletion in *Gm27162*^-/-^ mice is denoted by orange vertical bar and the 4.2 kbp deletion in CTCF binding site^-/-^ is denoted by blue vertical bar. **(e-g)** Heatmaps display the pairwise contact frequency maps measured by HiC in wildtype, *Gm27162^-/-^* and CTCF binding site^-/-^ double positive (DP) T cells using 30kbp resolution.

Among paralogous TF pairs, we focused on the ETS family proteins because of their conservation in metazoans over a large span of evolutionary time and their roles in regulating hematopoiesis and immune responses^13^. There are four ETS pairs as defined above in both human and mouse genomes: *ETS1* and *FLI1*, *ETS2* and *ERG*, *ELF5* and *EHF*, in addition to *ETV3* and *ETV3L* (Figure 1c). We found *ETS1-FLI1, ETS2-ERG* and *ELF5-EHF* pairs to be within 100-300 kbp of each other in an anti-sense orientation. In contrast, the *ETV3L–ETV3* pair is positioned more proximally, approximately 30 kb apart, with both genes transcribed from the same strand. Using publicly available single-cell RNA-seq data^28, 29^, we analyzed the expression patterns of these TFs across immune cell subsets in human lymph nodes (Figure S1a). *ETS1* and *FLI1* are highly expressed in T and B lymphocytes, and NK cells, with *ETS1* consistently exhibiting higher expression levels than *FLI1* in T and NK cells (Figure S1a). In myeloid cells, *ETS2* and *FLI1* display high transcriptional activity. In contrast, the *ELF5-EHF* and *ETV3L-ETV3* pairs showed low to moderate transcriptional activity across immune cell types (Figure S1a). Based on the strong expression of the *ETS1-FLI1* pair in T cells compared to other paralogous pairs and cell types, we aimed to study this paralogous TF pair in T cell contexts.

### Enhancer connectivity at the *Ets1-Fli1* locus is flanked by H3K27me3-rich repressed chromatin

Previously, we showed that the ∼1 Mbp DNA sequence harboring *Ets1* and *Fli1* genes forms a highly connected multi-enhancer hub with exceptional degree of connectivity in T cells^19, 20^. This locus has a unique histone post-translational modification structure with stretches of highly acetylated segments spanning the *Ets1* and *Fli1* promoters and gene-bodies in addition to a 25kbp segment downstream of *Ets1*, which scores as a super-enhancer selectively in T cells. This super-enhancer, which we previously referred to as *Ets1*-SE^20^, is also annotated as a long non-coding RNA *Gm27162*. The orthologous region in humans harbors multiple single nucleotide polymorphisms (SNPs) implicated in allergy, asthma^30^ and atopic dermatitis^31^. Previously, we generated a mouse strain with a genetic deletion of *Gm27162* DNA sequence to understand the implications of genetic perturbation at this locus in T cell biology^20^. We reported that deletion of *Gm27162* was dispensable for T cell development. However, it impaired the differentiation of CD4^+^ T helper 1 (Th1) cells. This impairment produced an allergy phenotype in *Gm27162^-/-^* mice due to the failure of balancing Th1 and Th2 differentiation^20^. This prior work suggests genetic perturbation of the super-enhancer *Gm27162* does not affect T cell development but can change T cell function^20^. However, the *cis* regulation of chromatin fiber is unknown. We reasoned that a detailed understanding of how chromatin architecture is rewired following such genetic perturbations could provide insight into the conserved genomic positioning of these two paralogous TFs in vertebrates.

We first examined histone modifications at this locus. To complement our previously generated histone acetylation data, we assessed if the *Ets1-Fli1* locus is marked with repressed histone modification H3K27me3 by performing cleavage under targets & release using nuclease (CUT&RUN^32^) in double-positive (DP) CD4^+^CD8^+^ T cells collected from the thymus (Figure 1d). Enhancer connectivity measured by HiChIP was enriched within a stretch of H3K27ac modification, flanked by two H3K27me3 domains which are ∼750 kbp apart (Figure 1d). Hence, the paralogous TFs *Ets1* and *Fli1* are embedded within an ‘island’^33^ of active chromatin forming a multi-enhancer hub flanked by repressed chromatin. Intrigued by this separation of active and repressed histone modifications, we generated a novel mouse strain, referred to as CTCF binding site^-/-^, where we deleted a 4.2kbp fragment enriched for CTCF binding sites at the boundary between the H3K27ac-H3K27me3 transition downstream of the *Ets1* gene (blue bar, Figure 1d). Thus, *Gm27162^-/-^ and* CTCF binding site^-/-^ mouse strains are the genetic tools we used to study *cis* regulation and genome rewiring at the *Ets1-Fli1* locus.

We first performed HiC^34, 35^ in DP T cells from the mouse thymus (Figure 1e-g). Examination of HiC contact frequency map in wildtype T cells showed the establishment of three domain structures insulated by CTCF binding sites and significant interactions between *Ets1* and its super-enhancer, *Gm27162* (Figure 1e). The largest domain structure which encompasses *Ets1* and *Gm27162* is bordered by convergent CTCF binding motifs (Figure 1d, arrows represent CTCF motif orientation). The deletion of *Gm27162* led to fewer long-range interactions through less frequent contacts compared with wildtype T cells (Figures 1f, S1b). The deletion of the CTCF binding site at the boundary of H3K27ac and H3K27me3 domains led to loss of insulation between the two domains (Figures 1g, S1b). Together, the population-level HiC data suggested broad changes in chromatin organization at the *Ets1-Fli1* locus after deletion of the super-enhancer or the CTCF boundary element.

### Chromatin tracing reveals spatial proximity between flanking H3K27me3 domains

While population-level assays like HiC provide valuable insights into general chromatin architecture, they are fundamentally incapable of capturing the full spectrum of chromatin folding configurations for genes and their regulatory elements. This is because HiC and similar sequencing assays fragment the chromatin of millions of cells and cannot resolve allele-specific interactions. We implemented Optical Reconstruction of Chromatin Architecture (ORCA)^6, 24, 36–38^ to trace the chromatin fiber at the *Ets1-Fli1* locus in DP T cells. We designed the ORCA primary probes to tile a 900 kbp window at a resolution of 30 kbp for sequential labeling of this locus. We hybridized the probes on primary DP T cells and used microscopy to reconstruct the chromatin fiber at the single-allele level, collectively generating ∼10,000 chromatin traces (Figure 2a). Each 30 kbp segment was imaged using unique readout probes with high detection efficiency resolved into 30 (3D) coordinates, forming a sequentially labeled chromatin ‘walk’ (Figure S2a). A walk is a contiguous series of imaged loci that traces the spatial organization of an individual chromatin fiber. Our ORCA experiments were specifically optimized to improve the full-trace detection percentage to avoid imputation of missing readouts (Figure S2b). We took a highly stringent strategy and excluded chromatin traces that missed hybridization to any readout probes. To assess the quality of our imaging-based approach, we pooled chromatin traces measured by ORCA and mapped average pairwise distances which generated a similar pattern of domains, loops, and stripes as observed in the contact frequency map from HiC (R^2^ = 0.86, Figure 2b). We defined ‘interactions’ in our chromatin tracing measurements based on 3D distances falling below a stringent threshold (<150 nm) which has been proposed in previous studies^6, 37^. The ORCA contact frequency matrix based on thresholding 3D distances correlated highly with the HiC contact frequency matrix (R^2^ = 0.9, Figure 2c). We also reported strong reproducibility between ORCA experiments in two biological replicates (Figure S2c,d). Throughout this study, we performed ORCA experiments in two biological replicates and pooled thousands of individual chromatin traces across two replicates for downstream analysis. Thus, our ORCA-based chromatin tracing experiments closely recapitulate the chromatin architecture observed in population-level HiC assays, while adding high-resolution, single-allele insights into chromatin organization.

**Figure 2.**
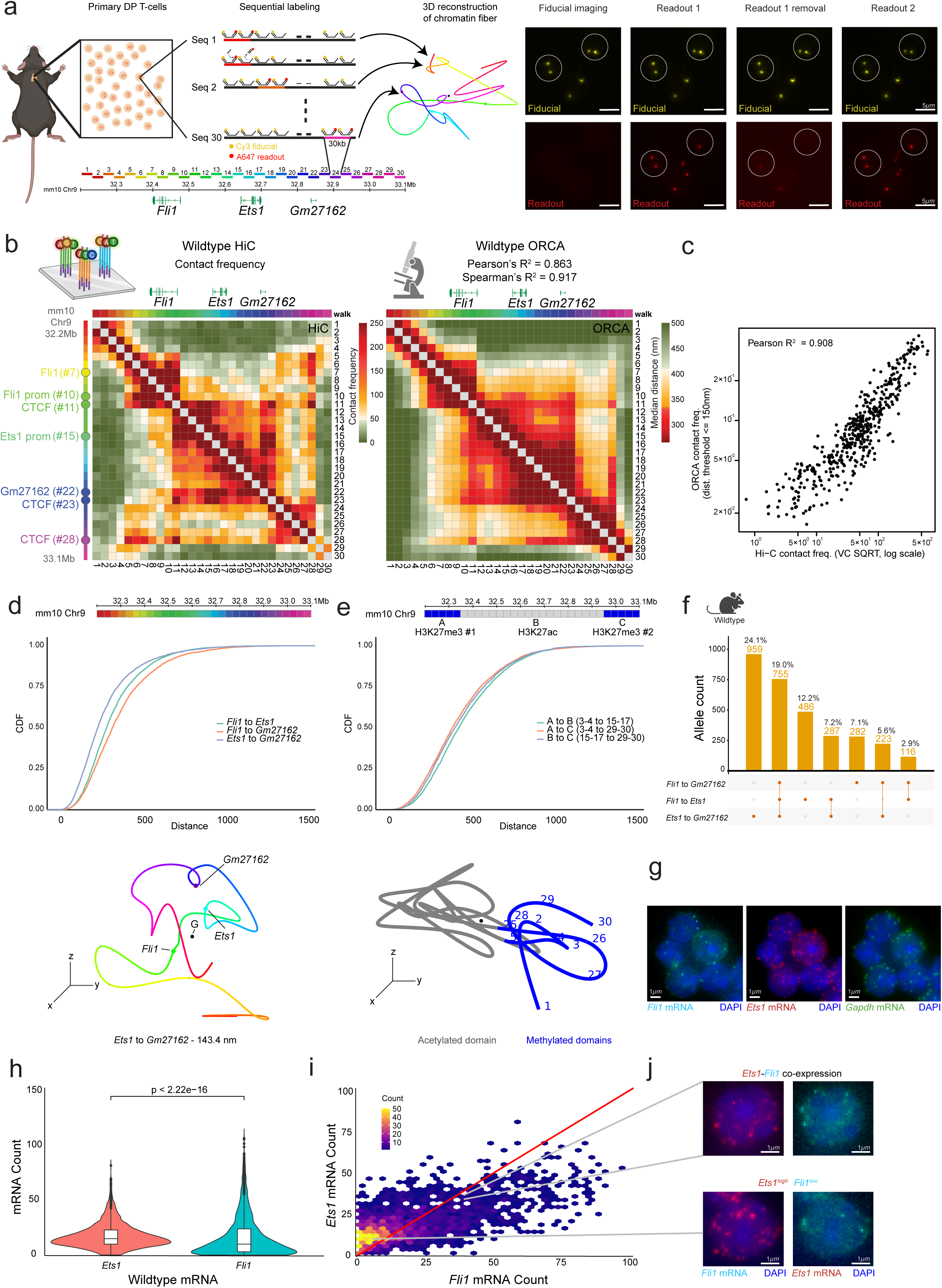
Chromatin tracing reveals spatial proximity between flanking H3K27me3 domains. **(a)** Schematic of ORCA primary probe tiling and chromatin trace reconstruction. CD4^+^ CD8^+^ double-positive (DP) T cells were extracted from the thymus, hybridized with ORCA primary probes and sequentially labeled with unique readout sequences. ORCA readouts unique for every 30 kbps at the *Ets1-Fli1* locus is pseudo-colored above a genome track annotated with the chromosome number and linear genome distance in 100 kbp increments. Readout probes were hybridized, imaged and displaced by a complementary toehold sequence before the introduction of the next 30 kbp readout sequence. Representative images show the ORCA labeling strategy (readout hybridization and toehold-dependent removal) using a reference (also called fiducial) and readout channel for imaging drift correction and sequential chromatin tracing respectively. **(b)** Comparison of wildtype HiC contact frequency map binned at 30 kbp resolution **(left)** with ORCA pairwise-distance matrix **(right).** Color bar represents the contact frequency and median distance for HiC and ORCA, respectively. Pearson’s and Spearman’s R^2^ values show the correlation between the HiC contact frequency matrix and ORCA median distance matrix. **(c)** Correlation between HiC and ORCA contact frequency displayed as Pearson’s R^2^ value (threshold used to define contact in ORCA = 150 nm). **(d) (Top)** Cumulative distance distribution between the centroids of *Fli1* to *Ets1* (green, mean = 332.0 nm), *Fli1* to *Gm27162* (orange, mean = 372.0 nm) and *Ets1* to *Gm27162* (blue, mean = 285.3 nm). **(Bottom)** Representative ORCA chromatin trace shows the proximity between *Ets1* and its super-enhancer *Gm27162* (*Ets1 – Gm27162* distance = 143.4 nm). Color bar shows the pseudo-color assigned to the 30 kbp readout sequences used to sequentially image the *Ets1-Fli1* locus. **(e) (Top)** Cumulative distance distribution between the centroids of acetylated and methylated chromatin regions grouped in Figure 1a, methylated domain #1 (A) to acetylated domain (B) (green, mean = 472.4 nm), methylated domain #1 (A) to methylated domain #2 (C) (orange, mean = 438.4 nm), acetylated domain (B) to methylated domain #2 (C) (blue, mean = 450.7 nm). **(Bottom)** Representative ORCA chromatin trace shows the association of methylated domains in 3D space despite the 750 kbp separation in 1D genomic distance. Color bar above plot represents the position of the two methylated domains (blue) and the acetylated domain (grey) on a 1D genome scale annotated with chromosome number and coordinates. **(f)** Upset plot showing the frequency and allele-number of three-way interactions between *Ets1, Fli1* and *Gm27162* (contact threshold = 257.89 nm, total chromatin traces analyzed = 3979) **(g)** Representative images of single molecule RNA FISH showing the labeling of *Fli1* mRNA (cyan), *Ets1* mRNA (red) and *Gapdh* mRNA (green) in DP T cells overlaid with DAPI nuclei staining (blue). **(h)** Violin plot of *Ets1* and *Fli1* mRNA count in wildtype DP T cells (significance computed using two-sided Wilcoxon rank sum test) **(i)** 2D histogram displaying the level of *Ets1* and *Fli1* mRNA recorded per wildtype DP T cell. Color bar shows the number of cells within each bin **(j)** Representative single molecule RNA FISH images showing *Ets1-Fli1* co-expressing cells **(top)** and *Ets1*^high^ *Fli1*^low^ cells **(bottom)** mapped to their respective areas within Figure 1i.

Key genomic regions within this locus, including the *Ets1* and *Fli1* gene bodies and their flanking H3K27me3-marked domains, extend beyond 30 kbp, spanning multiple readout probes in our chromatin tracing experiments. To evaluate the spatial proximity of regions of interest, we determined the geometric centers, also called centroids, spanning *Fli1* and *Ets1* gene loci in addition to *Gm27162* and calculated the Euclidean distances between the centroids per allele. Cumulative distance distribution analysis across single alleles for the centroids of *Ets1, Fli1,* and *Gm27162* confirmed that the *Ets1* gene is most frequently positioned near its super-enhancer *Gm27162* (Mean distance = 285.3 nm, Figure 2d top), as seen in the representative chromatin trace (Figure 2d bottom).

Surprisingly, the 3D distance between the two flanking H3K27me3-marked regions (Mean = 438.4 nm) was smaller compared with the 3D distance between regions with opposite histone modifications (Mean = 450.7 nm or 472.4 nm) despite a linear genomic distance of 750 kbps between the two H3K27me3 regions (Figure 2e top). A representative trace showcases the average trend (Figure 2e bottom). This spatial proximity was not detected in bulk HiC experiments (Figure 1e). Our microscopy technique which directly measures distances may overcome HiC’s limited enzymatic capture radius. Hence, the H3K27me3-rich domains could play a structural role in stabilizing the flanking regions of the *Ets1-Fli1* locus and promoting interactions within the central H3K27ac-rich multi-enhancer domain.

The *Ets1-Fli1* locus scored as a multi-enhancer hub in our bulk HiChIP analysis^19^. To quantify multi-way connectivity within this locus, we defined “interactions” based on 3D distances for individual alleles falling below a threshold. To avoid arbitrary thresholds, we used the average distance between consecutive segments as the interaction cutoff per experimental condition^37^. We detected high interaction frequencies between *Ets1*-*Gm27162* (24%) and to a lesser extent *Fli1-Ets1* (12%), indicating a competitive environment of interactions within the multi-enhancer hub (Figure 2f). This analysis also revealed that three-way interactions between *Ets1, Fli1,* and *Gm27162* regions were enriched in ∼19% of chromatin traces in wildtype DP T cells (Figure 2f). According to the HiC contact frequency map and HiChIP analysis^19^, CTCF binding sites frequently contact with highly acetylated elements including the *Ets1* promoter. Using ORCA, we observed that in ∼12% of our alleles, CTCF binding sites colocalized with H3K27ac-rich domains to facilitate multi-way interactions (Figure S2e). These 5-way interactions occurred at a frequency ∼5.5 times higher than those observed at the *Sox2* locus in mouse embryonic stem cells, as measured by publicly available distance-matched ORCA data^37^ (Figure S2e). Although in DP T cells, the most frequent topology relates to spatial localization of *Ets1* and *Gm27162* and their separation from *Fli1*, multi-way interactions of *Ets1*, *Gm27162* and *Fli1* is also detected in a percentage of alleles. These multi-way interactions and the spatial proximity of *Ets1* and *Fli1* gene loci supports border bypassing and the formation of the stacked boundary conformation^25^ in a percentage of alleles.

Next, we performed single-molecule RNA FISH^39, 40^ to quantify the expression of *Ets1* and *Fli1* in DP T cells (Figure 2g-j). We measured *Gapdh* mRNA abundance as a positive control to validate mRNA staining quality and found no significant differences in *Gapdh* mRNA detection across multiple fields of view imaged with co-labeling of *Ets1* and *Fli1* mRNA in the same cells (Figure S2f). On average, *Ets1* mRNA molecules were 50% more abundant than *Fli1* mRNA molecules (Median=15 molecules for *Ets1* and 10 molecules for *Fli1*) (Figure 2h). We mapped *Ets1* and *Fli1* transcript dynamics within the same cell and observed an enrichment of cells with high *Ets1* mRNA molecules and low *Fli1* mRNA molecules, referring to them as *Ets1*^high^*Fli1*^low^ cells. However, we also detected a small subset of cells that co-expressed *Ets1* and *Fli1* at comparable levels, corresponding to cells grouped closer to the red line (Figure 2i). Examination of single molecule RNA FISH images grouped close to the red line showed more *Fli1* mRNA spots in cells co-expressing *Fli1* and *Ets1* in comparison with *Ets1*^high^*Fli1*^low^ cells (Figure 2j). The co-expression of *Ets1* and *Fli1* in T cells may result from border bypassing events including the spatial proximity of the highly acetylated genomic regions of *Ets1* and *Fli1* gene loci, observed in approximately 12.2% of chromatin traces or the formation of the *Ets1-Gm27162-Fli1* hub detected in around 19% of chromatin traces (Figure 2f). Representative ORCA chromatin traces support the *Ets1-Gm27162-Fli1* hub and *Ets1-Fli1* interaction conformations (Figure S2g). Taken together, the chromatin structures and transcriptional profiles at the *Ets1-Fli1* locus across thousands of wildtype DP T cells proposed a novel role for flanking H3K27me3-rich domains in stabilizing highly interacting acetylated chromatin regions. Moreover, both cooperative and competitive enhancer-promoter interaction states can take place between paralogous TF pairs within the same population of cells.

### Deletion of the *Ets1* super-enhancer *Gm27162* leads to spatial proximity of *Ets1* and *Fli1*

### promoters

We next assessed how the chromatin fiber rewires after genetic deletion of the *Gm27162* super-enhancer. We performed H3K27ac and H3K27me3 CUT&RUN in DP T cells of *Gm27162^-/-^* mice and found comparable histone modifications in T cells at this locus in two mouse strains (Figure 3a). To trace the *Ets1-Fli1* locus in *Gm27162*^-/-^ DP T cells, we hybridized the same set of primary probes used in wildtype cells, filtered and pooled the chromatin traces with 30 successful hybridizations (Figure S3a). We employed the 30 kbp hybridization scheme at the locus harboring *Gm27162* to specifically capture a 5 kbp region flanking the 25 kbp deletion. This design enabled us to profile the impact of super-enhancer loss by imaging the 5 kbp fragment which we call the *Gm27162* deletion flanking region (DFR) (Figure S3b). The contact frequency matrix based on pooled alleles measured by ORCA correlated strongly with HiC and the imaging results were highly reproducible between two biological replicates using both median distance and contact frequency measurements (Figures 3b,c and S3c,d).

**Figure 3.**
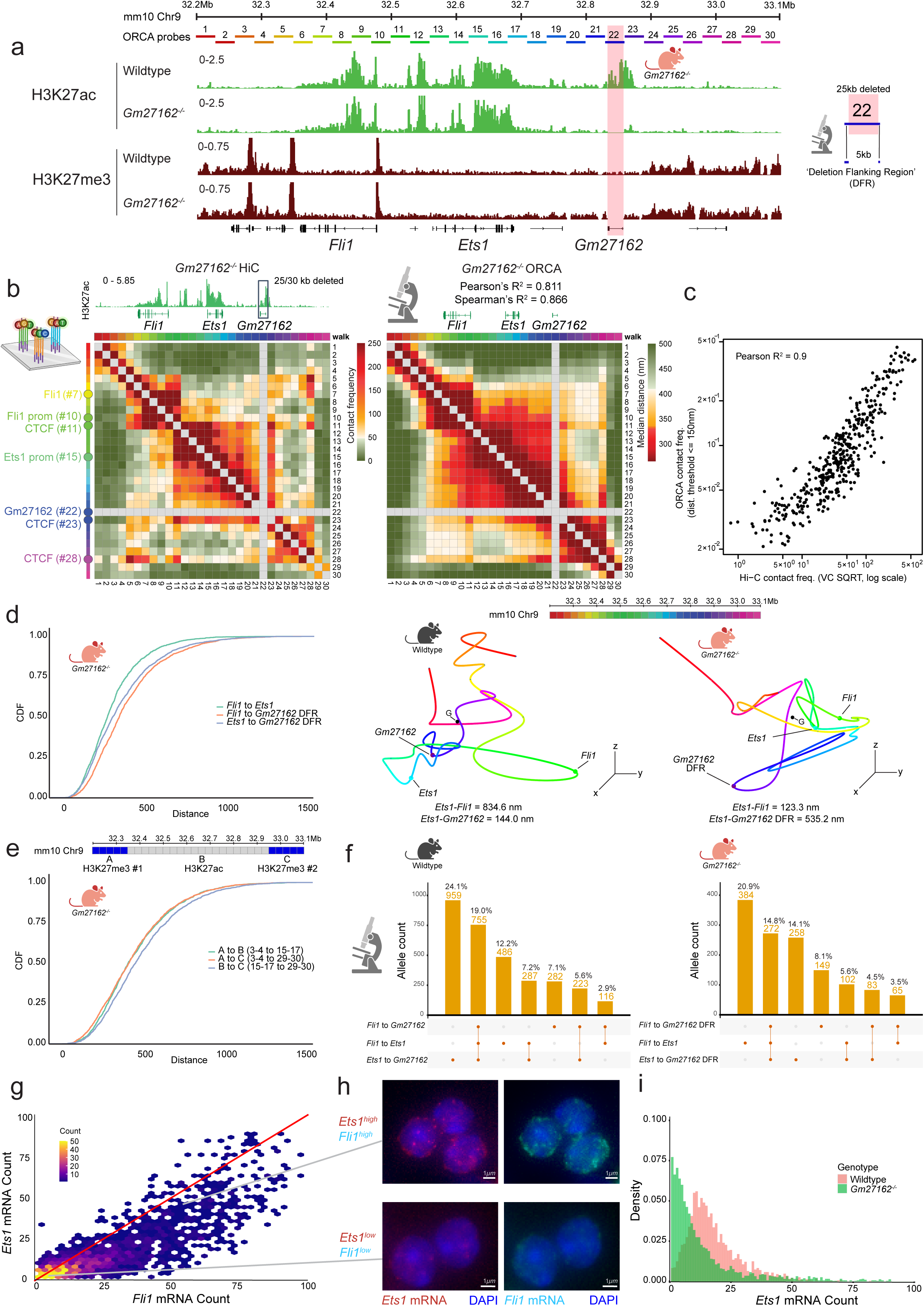
Deletion of the *Ets1* super-enhancer *Gm27162* leads to spatial proximity of *Ets1* and *Fli1* promoters. **(a)** Genome browser views comparing H3K27ac and H3K27me3 patterning within the *Ets1-Fli1* multi-enhancer hub measured by CUT&RUN in *Gm27162*^-/-^ DP T cells. Chromosome number, linear genome scale and 30 kbp ORCA readout segments are overlaid on CUT&RUN data. Orange bar represents the 25 kbp deletion of *Gm27162*. Schematic on the right shows that the 25 kbp deletion falls within a single 30 kbp ORCA bin (i.e., readout 22), with the remaining 5 kbp fragment defined as the *Gm27162* ‘Deletion Flanking Region’ or ‘DFR’. **(b)** Comparison of *Gm27162*^-/-^ HiC contact frequency map binned at 30 kbp resolution **(left)** with ORCA pairwise-distance matrix **(right)**. H3K27ac CUT&RUN genome browser track highlights the region deleted in the *Gm27162*^-/-^ mouse genotype. Color bar represents the contact frequency and median distance for the HiC and ORCA matrices, respectively. Pearson’s and Spearman’s R^2^ values show the correlation between the HiC contact frequency matrix and ORCA median distance matrix. **(c)** Correlation between *Gm27162*^-/-^ HiC and ORCA contact frequency displayed as Pearson’s R^2^ value (ORCA contact threshold = 150 nm). **(d)** Comparison of the cumulative distance distribution between the centroids of *Fli1* to *Ets1* (green, mean = 314.2 nm), *Fli1* to *Gm27162* (orange, mean = 402.4) and *Ets1* to *Gm27162* (blue, mean = 368.0 nm) in wildtype DP T cells **(left)**. Representative ORCA chromatin traces from *Gm27162*^-/-^ T cells show *Ets1-Fli1* promoter-promoter interactions **(right)** in contrast to wildtype where *Ets1-Gm27162* interactions are more frequent **(middle)**. Color bar shows the pseudo-color assigned to the 30 kbp readout sequences used to sequentially image the *Ets1-Fli1* locus. **(e)** Cumulative distance distribution between the centroids of acetylated and methylated chromatin regions in *Gm27162*^-/-^ DP T cells, methylated domain #1 (A) to acetylated domain (B) (green, mean = 467.5 nm), methylated domain #1 (A) to methylated domain #2 (C) (orange, mean = 459.7 nm), acetylated domain (B) to methylated domain #2 (C) (blue, mean = 506.1 nm). Color bar above plot represents the position of the two methylated domains (blue) and the acetylated domain (grey) on a 1D genome scale annotated with chromosome number and coordinates. **(f)** Upset plot comparing the changes in multi-way interaction frequency between wildtype DP T cells (contact threshold = 257.89 nm, total chromatin traces analyzed = 3979) **(left)** and *Gm27162*^-/-^ DP T cells (contact threshold = 260.99 nm, total chromatin traces analyzed = 1834) **(right)**. **(g)** 2D histogram plots comparing the ratio of *Ets1* and *Fli1* mRNA production per cell in *Gm27162*^-/-^ DP T cells. Color bar shows the number of cells within each bin. **(h)** Representative single molecule RNA FISH images showing *Ets1*^high^ *Fli1*^high^ cells **(top)** and *Ets1*^low^ *Fli1*^low^ cells (**bottom)** cropped from the same field of view and mapped to their respective areas within Figure 3i DAPI staining is overlaid on *Ets1* and *Fli1* mRNA images showing nuclear area. **(i)** Density histogram displaying the level of *Ets1* mRNA production in wildtype DP T cells (orange) and *Gm27162*^-/-^ DP T cells (green).

Next, we computed the distribution of centroid distances between *Ets1, Fli1* gene loci and the *Gm27162* DFR to pinpoint the precise reorganization within the locus upon deletion of the super-enhancer. Surprisingly, the *Ets1* and *Fli1* gene-bodies were more proximal to each other in *Gm27162*^-/-^ compared to wildtype T cells (Mean=332 nm in wildtype and 314.2 nm in *Gm27162*^-/-^, Figure 3d). This increased proximity was also observed when focusing specifically on the walks corresponding to the *Ets1* and *Fli1* promoters (Figure S3e). Additionally, the *Ets1* gene locus was further away in 3D distance from the *Gm27162* DFR (Mean=285 nm in wildtype and 368 nm in *Gm27162*^-/-^, Figure 3d). This suggested to us that without the *Gm27162* super-enhancer, the 5kbp *Gm27162* DFR loses its propensity to move proximal to *Ets1.* Representative ORCA chromatin traces corroborate this observation (Figure 3d, right). Distance between H3K27me3 modified regions did not change in *Gm27162*^-/-^ compared with wildtype cells (Figure 3e). To assess the chromatin compactness of the region spanning *Fli1* and *Ets1*, we computed the radius of gyration (Rg), which is a measure of the spatial distribution of chromatin relative to its center of mass^41, 42^ (Figure S3f). We found an increased frequency of chromatin traces where the region spanning *Ets1* and *Fli1* was more compact in *Gm27162^-/-^*DP T cells compared to wildtype counterparts (Figure S3g). To further quantify this reorganization, we assessed the frequency of multi-way interactions between *Fli1, Ets1* and the *Gm27162* DFR (Figure 3f). We detected a reduction in forming three-way interactions (19% to 14%) and a reduction in *Ets1-Gm27162* interaction (24% to 14%) (Figure 3f). Contrasting this loss, we found an increase in chromatin traces demonstrating spatial proximity of *Fli1* and *Ets1* in *Gm27162^-/-^*DP T cells (12% to 20%) (Figure 3f). We next examined multi-way interactions facilitated by CTCF binding sites. We found a reduction in forming multi-way contacts in *Gm27162^-/-^* cells compared with wildtype cells, albeit the frequency was still higher than random chance as seen by the selection of distance matched readouts from the *Sox2* locus in mouse embryonic stem cells^37^ (Figure S3h). Representative chromatin traces in *Gm27162^-/-^* cells confirmed the reduced spatial proximity and increased association between *Fli1* and *Ets1* (Figure S3i). Together, loss of the super-enhancer *Gm27162* leads to spatial proximity of *Ets1* and *Fli1* gene loci which are insulated in distinct domains.

We next sought to evaluate the functional consequences of this spatial proximity between paralogs *Fli1* and *Ets1*. Using single molecule RNA FISH, we found a depletion of *Ets1*^high^*Fli1*^low^ cells (Figure 3g). Instead, there was an enrichment of cells with high levels of *Fli1* transcription (*Fli1*^high^) with two- to three-fold increase in *Ets1* production, marked by a higher frequency of cells close to the red line (Figure 3g). These RNA FISH data suggest co-expression of *Ets1* and *Fli1* in *Gm27162^-/-^* T cells (Figure 3g). Visual inspection of representative *Ets1*^high^ *Fli1*^high^ cells showed higher detectable *Ets1* and *Fli1* mRNA spots in comparison with *Ets1*^low^ *Fli1*^low^ cells selected from the same field of view imaged using *Gm27162^-/-^* DP T cells (Figure 3h). Despite the detection of this population, only a small fraction of cells produced *Ets1* transcript levels equivalent to or higher than the mean transcript level observed in wildtype DP T cells, indicated by the overlapping area in the histogram (Figure 3i). Thus, in the absence of the *Gm27162* super-enhancer in DP T cells, border bypassing^25^ and the interactions between *Fli1* and *Ets1* increases, leading to an increase in the proportion of cells co-expressing *Fli1* and *Ets1*.

### Deletion of the CTCF binding site at the boundary between H3K27ac-H3K27me3 regions causes domain-wide decompaction

We next sought to examine how deleting the CTCF binding sites separating the stretch of highly acetylated elements from the repressed regions with H3K27me3 modification rewires the chromatin architecture. Assessing changes in histone modification, we found that deletion of the CTCF binding sites led to the spread of H3K27ac into the previously H3K27me3-rich domain (dashed box, Figure 4a). The spread of H3K27ac was contained by another CTCF binding site ∼150 kbp downstream, which demonstrated an increase in CTCF occupancy and formed a strong boundary in CTCF binding site^-/-^ T cells (red bar, Figure 4a). Together, the chromatin state of the locus in CTCF binding site^-/-^ T cells downstream of the deletion shifts from a repressed to a partially repressed state.

**Figure 4.**
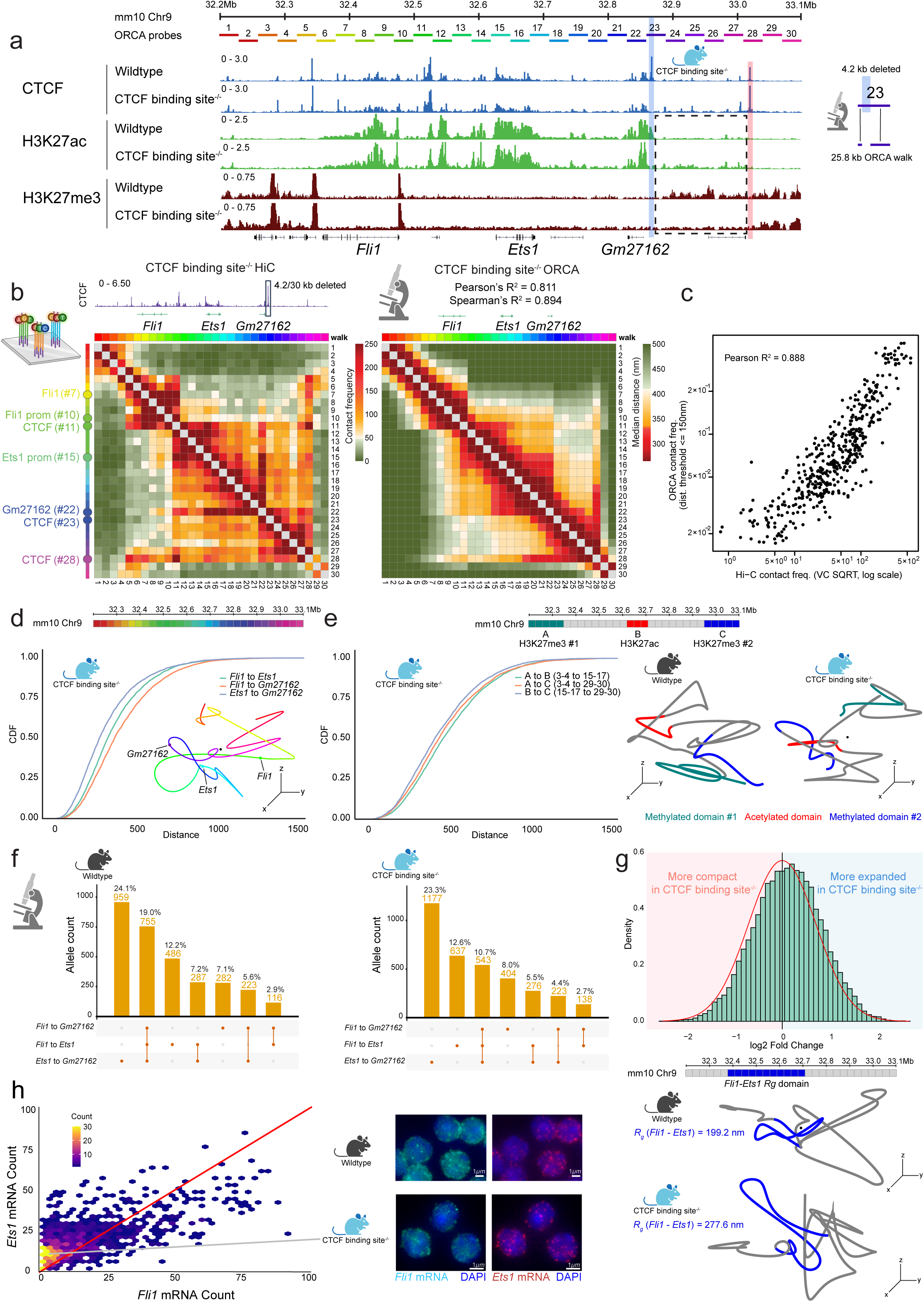
Deletion of the CTCF binding site at the interface between H3K27ac-H3K27me3 transition causes domain-wide decompaction at the *Ets1-Fli1* locus. (a) Genome browser views comparing H3K27ac and H3K27me3 patterning within the *Ets1-Fli1* multi-enhancer hub measured by CUT&RUN in CTCF binding site^-/-^ DP T cells. Changes in CTCF binding site occupancy between wildtype and CTCF binding site^-/-^ DP T cells are shown by CTCF CUT&RUN. Blue bar represents the 4.2 kbp CTCF binding site deletion at the interface between H3K27ac-H3K27me3 domains. Chromosome number, linear genome scale and 30 kbp ORCA readout segments are overlaid on CUT&RUN data. Schematic on the right shows that the 4.2 kbp deletion falls within a single 30 kbp ORCA bin (i.e., readout 23), with the remaining 25.8 kbp detected using ORCA imaging. Dashed rectangular box highlights the area of *de novo* H3K27ac spreading into H3K27me3 domain which is restricted from further invasion by CTCF binding site downstream (red). (b) Comparison of CTCF binding site^-/-^ HiC contact frequency map binned at 30 kbp resolution **(left)** with ORCA pairwise-distance matrix **(right)**. CTCF CUT&RUN genome browser track illustrates the proportion of ORCA bin deleted which corresponds to the CTCF binding site deletion in CTCF binding site^-/-^ mice. Color bar represents the contact frequency and median distance for the HiC and ORCA matrices, respectively. Pearson’s and Spearman’s R^2^ values show the correlation between the HiC contact frequency and ORCA median distance matrices. (c) Correlation between CTCF binding site^-/-^ HiC and ORCA contact frequency displayed as Pearson’s R^2^ value (ORCA contact threshold = 150 nm). (d) Cumulative distance distribution between the centroids of *Fli1* to *Ets1* (green, mean = 383.5 nm), *Fli1* to *Gm27162* (orange, mean = 427.2 nm) and *Ets1* to *Gm27162* (blue, mean = 338.8 nm) in CTCF binding site^-/-^ DP T cells. Representative ORCA chromatin trace shows proximity between *Ets1-Gm27162* elements (3D distance = 245.1 nm) and increased separation between *Ets1-Fli1* (3D distance = 418.2 nm). Color bar shows the pseudo-color assigned to the 30 kbp readout sequences used to sequentially image the *Ets1-Fli1* locus. (e) Comparison of cumulative distance distribution between the centroids of acetylated and methylated regions in CTCF binding site^-/-^ DP T cells **(left)**, methylated domain #1 (A) to acetylated domain (B) (green, mean = 511.6 nm), methylated domain #1 (A) to methylated domain #2 (C) (orange, 493.9 nm), acetylated domain (B) to methylated domain #2 (C) (blue, 464.7 nm). Color bar above plot represents the position of methylated domain #1 (green), acetylated domain (red) and methylated domain #2 (blue) on a 1D genome scale annotated with chromosome number and coordinates. Representative ORCA chromatin traces show increased association between the *de novo* H3K27ac domain with the centrally positioned H3K27ac domain in CTCF binding site^-/-^ mice **(right)**, compared to wildtype DP T cells which have both H3K27me3 domains in proximity **(middle)**. (f) Upset plot comparing the changes in multi-way interaction frequency between wildtype DP T cells (contact threshold = 257.89 nm, total chromatin traces analyzed = 3979) **(left)** and CTCF binding site^-/-^ DP T cells (contact threshold = 260.99 nm, total chromatin traces analyzed = 5059) **(right)**. (g) Histogram of log2(FC) local radius of gyration (Rg) ratio between CTCF binding site^-/-^ DP T cells and wildtype DP T cells **(top)**. The y-axis represents the frequency of observing a particular log fold change which was computed by bootstrapping 1000 alleles per genotype for 1000 iterations and calculating the local Rg ratio at each iteration to account for differences in total allele count between CTCF binding site^-/-^ DP T cells and wildtype DP T cells. The red area corresponds to more compact chromatin structure whereas the blue area corresponds to a more expanded chromatin structure. The red curve represents the shape of the distribution if it was normally distributed. Representative ORCA chromatin traces from wildtype **(middle)** and CTCF binding site^-/-^ DP T cells (**bottom)** show the increase in radius of gyration observed between the regions spanning *Fli1* and *Ets1*. (h) 2D histogram plots comparing *Ets1* and *Fli1* mRNA per cell in CTCF binding site^-/-^ DP T cells **(left)**. Color bar shows the number of cells within each bin. Representative single molecule RNA FISH images for wildtype **(right, top)** and CTCF binding site^-/-^ DP T cells **(right, bottom)** show the decrease in *Fli1* mRNA spots detected in CTCF binding site^-/-^ DP T cells. Arrow represents the area in Figure 4h which correspond to the representative image selected. DAPI staining is overlaid on *Ets1* and *Fli1* mRNA images showing nuclear area.

To uncover chromatin rewiring at this locus following the disruption of the H3K27ac-H3K27me3 patterning, we next performed chromatin tracing experiments on CTCF binding site^-/-^ DP T cells. When we filtered and pooled chromatin traces with all 30 readouts hybridized, chromatin traces measured by ORCA correlated strongly with HiC (Figures 4b-c and S4a). We also detected a strong correlation between two biological replicates using both median distance and contact frequency measurements (Figure S4b-c). With single cell chromatin traces available, we asked how the distribution of centroid distances between *Ets1, Fli1* and *Gm27162* changed in CTCF binding site^-/-^ T cells. Similar to wildtype T cells, *Ets1* and *Gm27162* were the most proximal interacting pair in 3D space in CTCF binding site^-/-^ T cells (Figure 4d). A representative ORCA chromatin trace shows the maintenance of *Ets1-Gm27162* spatial proximity (Figure 4d). Surprisingly, the mean distance distribution between different genomic regions was bigger by more than 50 nm in CTCF binding site^-/-^ DP T cells relative to wildtype counterparts (Figure 4d). Next, we wondered if spreading of H3K27ac into the H3K27me3 region downstream of the CTCF binding site deletion would impact the spatial localization of H3K27me3 domains. Measuring the distances between the centroids of H3K27ac and H3K27me3 domains revealed a larger distance between the flanking regions in CTCF binding site^-/-^ T cells compared with wildtype T cells (Mean=438nm in wildtype and 493nm in CTCF binding site^-/-^, Figure 4e). Representative ORCA chromatin traces show the intermixing between H3K27ac and the *de novo* H3K27ac chromatin domains which contrasts the association of H3K27me3 flanking domains in wildtype DP T cells (Figure 4e). These data suggest the CTCF binding sites and the histone modification state tightly control chromatin organization of this locus.

We next computed three-way interactions between *Ets1, Fli1* and *Gm27162* and reported a domain-wide reduction in multi-way interactions, particularly all elements coming together decreased after CTCF binding site deletion (19% to 10%, Figure 4f). Informed by HiC, we examined multi-way interactions facilitated by CTCF binding sites which revealed a reduction in forming multi-way contacts in CTCF binding site^-/-^ DP T cells (Figure S4d, 12.4% to 5.4%). We computed the local radius of gyration which revealed a higher frequency of traces had de-compacted or more expanded chromatin conformations (Figure S4e). These data support our observation of increased mean distance between interacting elements such as *Ets1* and *Fli1* (Figure 4d). This domain decompaction is seen as far as ∼400 kbp upstream of the CTCF binding site deletion between *Fli1* and *Ets1*, with a larger proportion of chromatin traces having a greater log fold change in radius of gyration relative to wildtype (Figure 4g). Representative ORCA chromatin traces visually support an expanded chromatin domain between *Fli1* and *Ets1* in CTCF binding site^-/-^ DP T cells relative to wildtype DP T cells (Figure 4g).

With domain-wide decompaction observed, we speculated whether *Ets1* and *Fli1* mRNA production was also impaired. Single molecule RNA FISH did not detect any significant change in *Ets1* production in line with the observation of conserved *Ets1-Gm27162* interactions (Figure 4h). However, we found a contraction in the proportion of *Fli1*^high^ cells, marked by a significant reduction in average *Fli1* mRNA expression (Figure S4f). Single molecule RNA FISH images confirm reduced *Fli1* mRNA spots while displaying comparable levels of *Ets1* mRNA relative to wildtype DP T cells (Figure 4h). Our chromatin tracing and single molecule RNA FISH data suggest that the *Ets1-Fli1* multi-enhancer hub undergoes a domain-wide decompaction in CTCF binding site^-/-^ T cells which reduces the proportion of chromatin traces engaging in long-range interactions driving *Fli1* expression in DP T cells (Figure S4g).

### Comparison of chromatin features in wildtype, *Gm27162*^-/-^ and CTCF binding site^-/-^ T cells

We next assessed additional features of chromatin organization by comparing wildtype, *Gm27162^-/-^*, and CTCF binding site^-/-^ T cells. We analyzed the organization of CTCF binding sites within chromatin traces, hypothesizing from previous studies^37, 43^ that the radial organization of CTCF binding sites is essential to coordinate interactions between *Fli1, Ets1* and *Gm27162* within the central acetylated domain. Using a distance-matched sliding window, we computed the frequency of interactions between three CTCF binding sites at boundaries within the *Ets1-Fli1* locus, situated within readouts #11, #23 and #28 in our ORCA probe design scheme (Figure 5a). We found that wildtype T cells had the highest frequency of spatial clustering of CTCF binding sites followed by *Gm27162^-/-^* cells (Figure 5a). The “centrality” analysis^37^ which accounts for the mean position of each readout plotted relative to the geometric center of each trace corroborated this observation, suggesting CTCF binding sites cluster around the geometric center (Figure S5a). As expected, CTCF binding site^-/-^ DP T cells demonstrated the lowest frequency of spatial proximity for CTCF bound boundaries. Moving the sliding window upstream, we observed interaction frequencies for distance-matched genomic regions excluding CTCF binding sites were also the lowest between the windows spanning *Fli1-Ets1* in in CTCF binding site^-/-^, corroborating our previous finding using local radius of gyration (Figure S5b). Representative ORCA chromatin traces show the radial organization of the CTCF boundary elements clustered around the geometric center in wildtype DP T cells which does not form in CTCF binding site^-/-^ DP T cells due to the deletion of CTCF binding site at readout #23 (Figure 5a).

**Figure 5.**
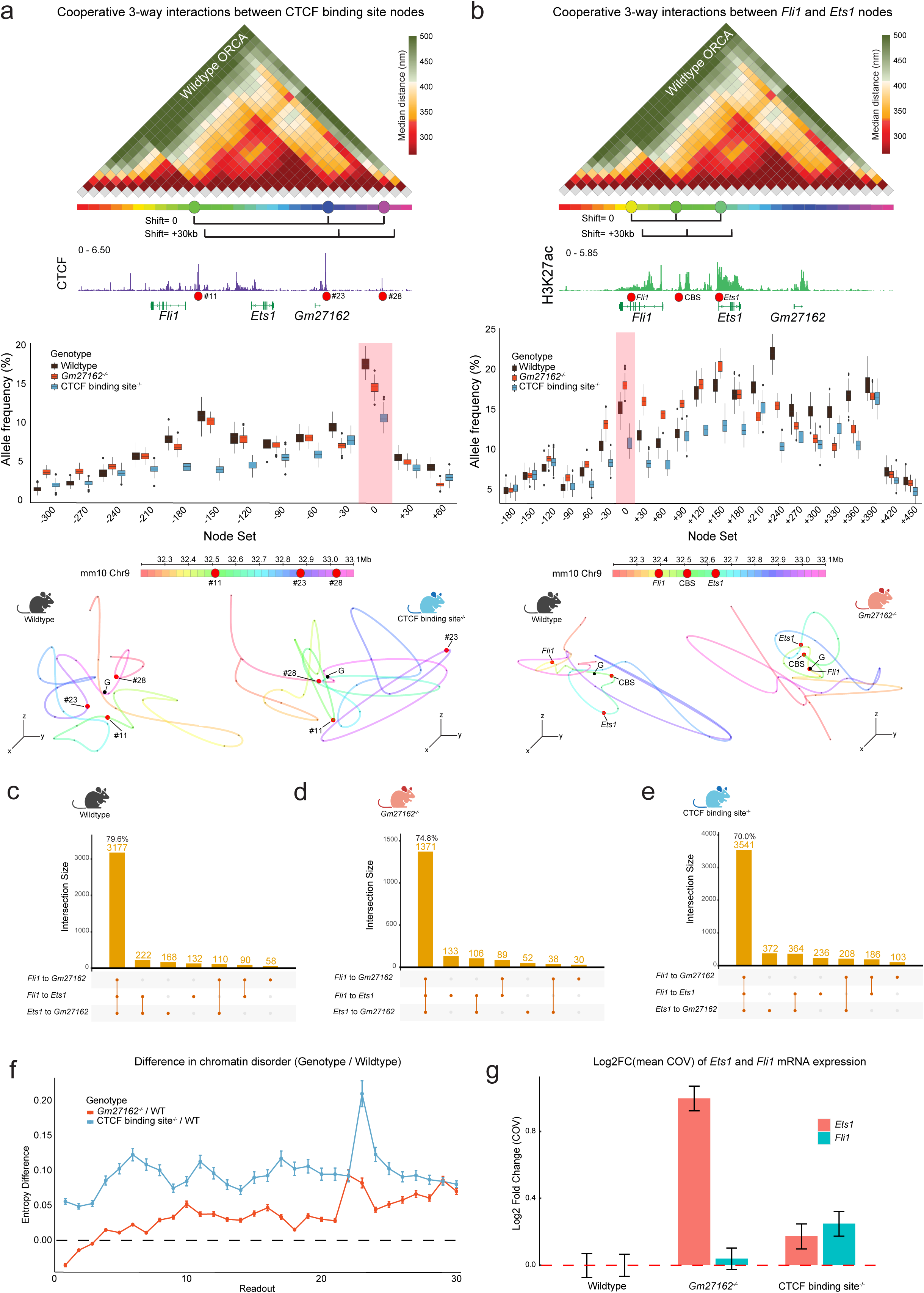
Comparison of chromatin features in wildtype, *Gm27162*^-/-^ and CTCF binding site^-/-^ T cells. (a) (Top) Wildtype ORCA pairwise distance map shows the *Ets1-Fli1* domain demarcated into 3 TAD domains based on the CTCF CUT&RUN genome browser track. Line segments represent the 3-way interaction window chosen. CTCF CUT&RUN genome track is also labeled with the ORCA readouts corresponding to the CTCF binding sites at the *Ets1-Fli1* locus which is used as the reference window for the 3-way interaction. **(Middle)** Boxplot showing the cooperative 3-way interaction frequency between CTCF binding sites in wildtype DP T cells (black), *Gm27162*^-/-^ DP T cells (red) and CTCF binding site^-/-^ DP T cells (blue). The y-axis represents the frequency of cooperative interactions computed by bootstrapping 1000 alleles per genotype for 1000 iterations. The number of interacting alleles was calculated per each iteration to account for differences in total allele count between the three genotypes. Interaction frequency of CTCF binding sites is highlighted by the red bar. Interaction frequency of distance matched non-CTCF binding site triplets along the *Ets1-Fli1* locus are shown as a function of distance (in kb) from the initial 3-way interaction window. **(Bottom)** Representative ORCA chromatin traces showing the radial organization of CTCF binding sites in wildtype DP T cells (left) and the perturbed radial organization in CTCF binding site^-/-^ DP T cells (right). Color bar shows the pseudo-color assigned to the 30 kbp readout sequences used to sequentially image the *Ets1-Fli1* locus along with the position of the CTCF binding sites examined in red. **(b) (Top)** H3K27ac genome browser track overlaid on wildtype ORCA pairwise distance map. Line segments represent the 3-way interaction window chosen between H3K27ac-rich regions. **(Middle)** Boxplot showing the cooperative 3-way interaction frequency between *Fli1*, *Ets1* and CTCF binding site (CBS) between the two genes based on high H3K27ac signal in wildtype DP T cells (black), *Gm27162*^-/-^ DP T cells (red) and CTCF binding site^-/-^ DP T cells (blue). The y-axis represents the frequency of cooperative interactions which was computed by bootstrapping 1000 alleles per genotype for 1000 iterations and calculating the number of interacting alleles each iteration to account for differences in total allele count between the three genotypes. Red bar shows the interaction frequency between *Fli1, Ets1* and CBS which is enriched in *Gm27162*^-/-^ DP T cells. Interaction frequency of distance matched non-H3K27ac rich triplets along the *Ets1-Fli1* locus are shown as a function of distance (in kb) from the initial 3-way interaction window. **(Bottom)** Representative ORCA chromatin traces showing decreased distances between *Fli1, Ets1* and CBS in *Gm27162*^-/-^ DP T cells (right) compared to wildtype DP T cells (left) **(c-e)** Upset plot shows the multi-way interaction frequency between the *Fli1*, *Ets1* and *Gm27162* using a ‘community’ distance threshold defining interaction (threshold = 600 nm) in wildtype DP T cells **(c)**, *Gm27162*^-/-^ DP T cells **(e)** and CTCF binding site^-/-^ DP T cells **(e)** (f) Entropy difference computed per readout between *Gm27162*^-/-^ DP T cells (red) and CTCF binding site^-/-^ DP T cells (blue) relative to wildtype DP T cells. Error bars show 95% confidence intervals. (g) Bar plot shows the log2(fold change) in the coefficient of variation (COV) of *Ets1* mRNA (red) and *Fli1* mRNA (blue) expression. Error bars show 95% confidence intervals.

To quantify border bypassing^25^ in *Gm27162*^-/-^ T cells, we evaluated the spatial localization of regions with high H3K27ac signature in addition to a CTCF binding site (CBS) at the boundary of *Ets1* and *Fli1*. Using the distance-matched sliding window approach, we found an increase in *Ets1* and *Fli1* interactions in *Gm27162*^-/-^ T cells anchored around ORCA readout probes annotated as H3K27ac-rich and the boundary between *Ets1* and *Fli1* (Figure 5b, 10% to 20%). We also report an increased interaction in *Gm27162*^-/-^ T cells up to 120 kbp upstream and downstream of the initial acetylation window, indicating more interaction between the highly acetylated segments of chromatin spanning *Ets1* and *Fli1* in *Gm27162*^-/-^ DP T cells relative to other genotypes. Representative ORCA chromatin traces show greater spatial proximity between the *Fli1* promoter, the boundary between *Ets1* and *Fli1,* and *Ets1* promoter in *Gm27162*^-/-^ DP T cells relative to wildtype DP T cells (Figure 5b).

We also sought to understand whether regulatory elements such as CTCF and H3K27ac-rich chromatin positioned *Ets1* and *Fli1* into higher-order structures within the nucleus. A recent study^38^ had proposed super-enhancer elements tend to associate within ‘communities’ under the effects of RNA Polymerase II (Pol II) clustering and nuclear compartmentalization^38^. Using our ORCA measurements of three annotated super-enhancers at *Fli1, Ets1* and *Gm27162* regions, we speculated whether the ability to form ‘communities’ was impaired by our perturbations within the locus. We found that ∼79% of wildtype T cells had the three elements interacting at a community threshold of 600 nm defined in the previous study^38^ (Figure 5c). We detected 74.8% and 70% of chromatin traces that form super-enhancer communities in *Gm27162^-/-^*and CTCF binding site^-/-^ T cells, respectively (Figure 5d-e). Our data suggest that perturbation of *cis-* regulatory elements within the *Ets1-Fli1* paralogous TF hub does not dramatically impair the ability to form super-enhancer communities since the effect of the perturbation affects local 3D organization within the ‘communities’ rather than global positioning within the nucleus.

### Genetic perturbation increases domain-wide disorder in chromatin folding

We next assessed the stability of chromatin conformations in DP T cells after genetic perturbations. We computed entropy^37^, which is defined as the degree of disorder or variability in chromatin folding. Higher entropy indicates greater structural instability and increased heterogeneity in chromatin interactions, whereas lower entropy suggests a more stable and predictable genome organization. We found that both *Gm27162^-/-^* and CTCF binding site^-/-^ T cells had more disordered chromatin structure compared with wildtype T cells, suggesting the perturbed chromatin conformations were more heterogeneous and potentially transient (Figure 5f). Increased disorder in chromatin folding because of CTCF binding site deletion has been reported at the *Sox2* and *Hox* clusters^37^. Next, we assessed whether the increased disorder in chromatin structure could explain the differences in transcriptional outputs of *Ets1* and *Fli1*. We computed the coefficient of variation in gene expression inferred from single molecule RNA FISH and plotted the log fold change over wildtype levels for both genes. We found that *Ets1* levels were the most variable in *Gm27162^-/-^* T cells, suggesting that the compensatory effect provided by the *Fli1* promoter is less stable than the regulation provided by the cognate super-enhancer (Figure 5g). Positive log fold change in the *Fli1* coefficient of variation supported our chromatin-level observations of increased *Fli1-Ets1* interactions driving the cooperative expression of both genes. For the CTCF binding site^-/-^ T cells, we observed a positive log fold change for both *Ets1* and *Fli1* mRNA levels (Figure 5g). Overall, our microscopy measurements in one mature T cell type, DP T cells, suggest that the epigenetic state and the 3D chromatin organization of the paralogous *Ets1-Fli1* locus are highly sensitive to genetic perturbation, with disruptions in *Gm27162* and CTCF binding site leading to increased chromatin disorder, transcriptional variability, and altered cooperative gene regulation.

## Discussion

Beyond elucidating the *cis*-regulatory mechanisms governing *Ets1* and *Fli1*, our study provides broader insights into how chromatin topology influences paralogous TF function in immune cells. The competitive and cooperative interactions between *Ets1, Fli1*, and their associated regulatory elements resemble principles observed in other paralogous TF clusters, such as the *HOXA* genes^8^, where chromatin conformation dictates lineage-specific gene expression. Our results suggest that paralogous TFs, particularly those retained in close genomic neighborhoods, rely on higher-order chromatin structure to coordinate co-regulation which also serves as a buffer that protects TF expression in the event of genetic mutation. Such genomic arrangements can confer an evolutionary advantage to paralogous TFs with critical roles in driving development such as vertebrae and limb development^44–46^. Furthermore, the increased disorder in chromatin folding and transcriptional output observed upon perturbation of histone modifications and CTCF binding highlights the importance of stable chromatin architecture in fine-tuning TF expression and gene dosage control. Given the enrichment of disease-associated SNPs within this locus, our findings raise intriguing possibilities about how genetic and epigenetic variations affecting paralogous TF hubs contribute to immune-related diseases. Future studies targeting chromatin-based regulation of paralogous TFs may provide new therapeutic strategies for modulating immune cell function in disease contexts.

Our CTCF binding site^-/-^ dataset revisits a paradox of nuclear organization – HiC contact frequency showed increased cross-TAD interactions when the CTCF binding site was removed, but ORCA suggested domain expansion. We speculate that HiC and ORCA are revealing two distinctive but complementary views on chromatin reorganization when a CTCF boundary element is removed. Without the CTCF-mediated insulation, HiC captures the new cross-domain contacts as chromatin regions previously separated interact more freely. Our ORCA data suggests that removal of a CTCF boundary allows the chromatin to adopt a more relaxed, extended conformation. As HiC reflects contact probability, we reconcile both modalities by considering expanded chromatin domains to contain a larger interaction space that can form different conformations at the time of cell fixation. Mechanistically, we hypothesize that loop extrusion through the dynamic activity of cohesin could help mediate these conformations by sliding past the former boundary, forming more expanded chromatin loops. The increase in CTCF occupancy downstream of deleted CTCF binding site supports this interpretation, suggesting that the extruded DNA sequence harboring *Ets1* and *Gm27162* is a larger DNA segment in CTCF binding site^-/-^ T cells. Our observations based on HiC data can then be explained by the aggregation of hundreds of thousands of loop extrusion combinations creating more contact between the two TADs despite a more expanded chromatin domain. Similar observations reconciling HiC and other imaging modalities have been reported experimentally^37^ as well as through polymer simulations^47, 48^.

Compensatory responses repurposing enhancer-promoter interactions have been previously reported in *Mesp1-Mesp2* expression control in embryonic stem cells^49^, where the *Mesp2-* enhancer interacts more with the *Mesp1* promoter and increases its expression upon the loss of *Mesp2*. In our perturbation of the *Gm27162* super-enhancer, we find that the *Fli1* promoter forms increased interactions with the *Ets1* promoter acting as a *de novo* enhancer or ‘E-promoter’, an observation reported by genome-wide CRISPR screens^50^ and within IFN-alpha induced genes in the immune context of LPS-stimulated macrophages^51^. While the compensatory effect does not confer phenotypic robustness in our *Gm27162^-/-^* mouse genotype^20^, we suspect that the partial rescue of *Ets1* levels facilitated by the *Fli1* promoter is adequate to sustain T cell development and avoid severe proliferative defects seen in mice with homozygous deletion of *Ets1*^52–54^. This observation re-explains the advantage of positioning paralogous TF pairs in close proximity while pinpointing the reason for the evolutionary success of the ETS TF family which has established itself in multiple metazoan species from sponges to vertebrates^55, 56^. One question that remains to be answered is about the precise mechanism that drives *Fli1* promoter interactions with *Ets1.* Recent literature has implicated phase-separated structures harboring transcription factors and co-activator complexes to form around enhancer elements within 3D hubs^57^. We speculate co-activator complexes such as BRD4, p300 and Mediator subunits could facilitate the ‘E-promoter’ role by bridging the promoter of *Fli1* to *Ets1* due to high H3K27ac deposition on both these elements^58^. Since *Ets1* and *Fli1* are actively transcribed in T cells, another possibility explaining this 3D chromatin organization is the ability of RNA polymerases to serve as a barrier for cohesin loop extrusion^59, 60^. Under this hypothesis, we speculate that obstructed cohesin loop extrusion between the *Fli1* and *Ets1* promoters could reduce the 3D distance between both elements, thereby upregulating their gene expressions synergistically.

## Acknowledgments

The authors thank Vutara VXL Slack group in particular Guy Nir, Sarah Aufmkolk, and Ting Wu, in addition to Alistair Boettiger for helpful discussions and technical advice. We are grateful to Katherine Lupo for providing critical technical support. This work was supported by the Burroughs Welcome Fund, the Chan Zuckerberg Initiative Award, and NIH UC4 DK112217, U01 DK112217, R01AI168240, U01 DK127768, U01 DA052715 (G.V). and R01-CA-248041 and R01-CA-230800 (to R.B.F.).

## Supplementary figure Legends

**Figure S1.**
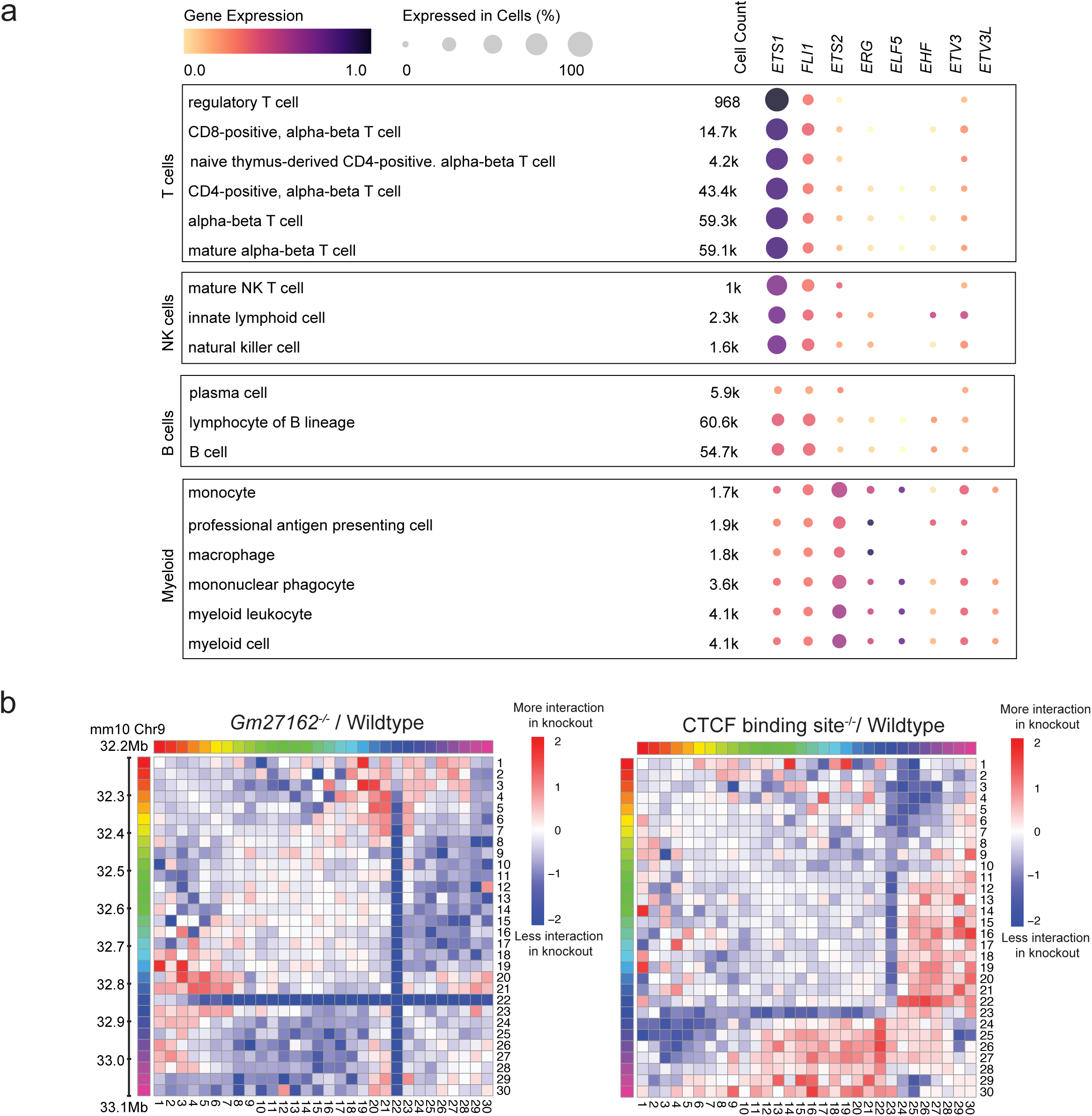
Enhancer connectivity at the *Ets1-Fli1* locus is flanked by H3K27me3-rich repressed chromatin. **(a)** Single cell RNA-seq data from The Tabula Sapiens available on cellxgene^28, 29^ showing the expression of paralogous TFs *ETS1, FLI1, ETS2, ERG, ELF5, EHF, ETV3* and *ETV3L* in immune cell populations such as T cells, NK cells, B cells and myeloid cells in human lymph nodes. Color bar represents the magnitude of TF expression. **(b) (Left)** HiC contact frequency map at 30 kb resolution showing the difference in interactions between *Gm27162*^-/-^ and wildtype DP T cells. **(Right)** HiC contact frequency map at 30 kb resolution showing the difference in interactions between CTCF binding site^-/-^ and wildtype DP T cells. Color bar indicates interaction state in the knockout mouse relative to wildtype – red represents more interactions in the knockout and blue represents less interaction between chromatin domains in the knockout mouse.

**Figure S2.**
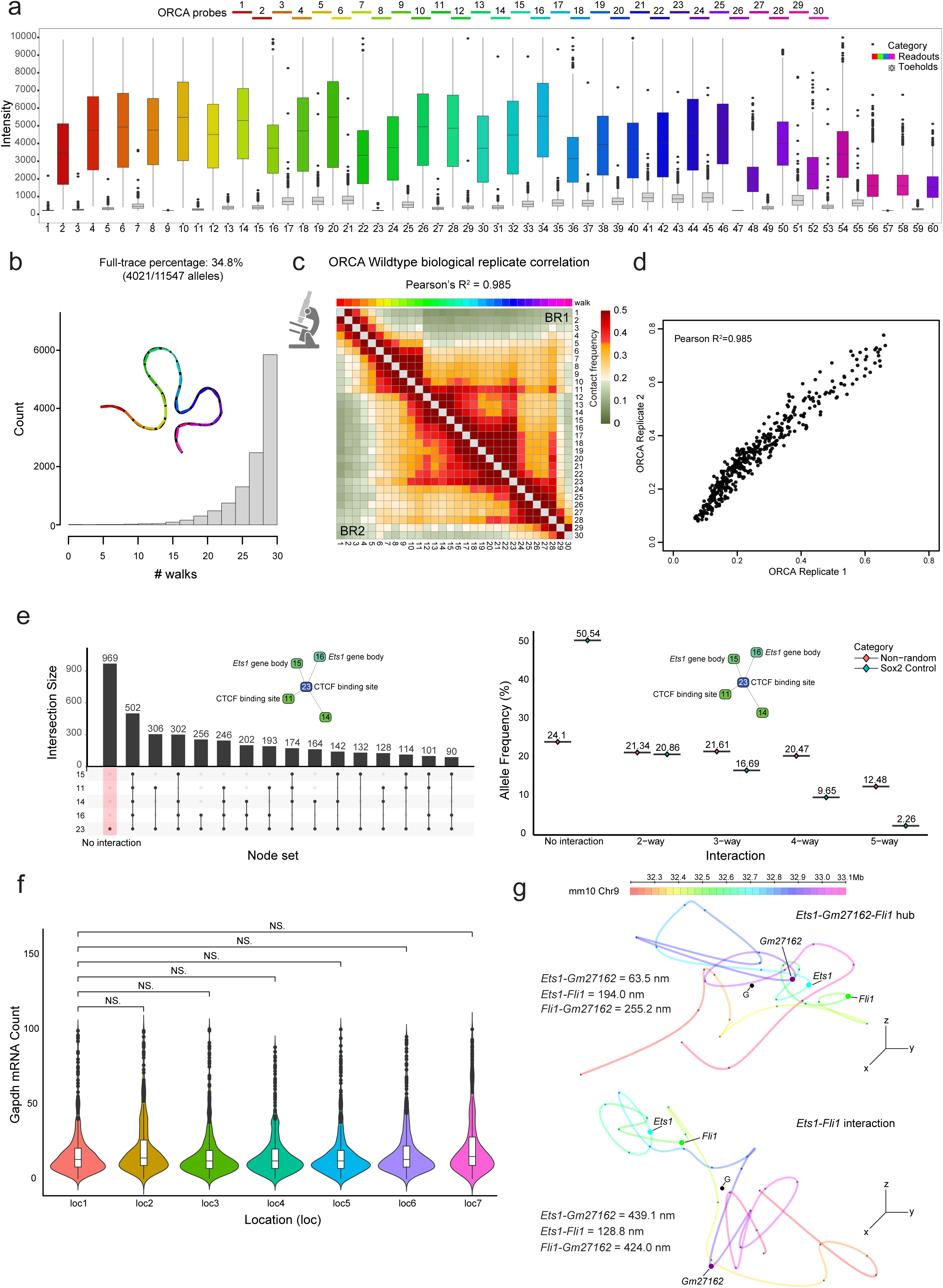
ORCA in wildtype T cells identifies spatial proximity between flanking H3K27me3 domains. **(a)** Boxplot of fluorescence signal intensity at each hybridization and readout removal/toehold step. Color bar shows the pseudo-color assigned to the 30 kbp readout sequences used to sequentially image the *Ets1-Fli1* locus. Grey boxplots represent toehold sequences and pseudo-colored boxplots correspond to readout hybridizations. **(b)** Bar plot displays the proportion of wildtype DP T cell chromatin traces as a function of the number of readouts successfully hybridized. Schematic of chromatin trace with all 30 successful hybridizations shown along with full-trace percentage. **(c)** Correlation between two ORCA wildtype biological replicates displayed as Pearson’s R^2^ value. **(d)** Comparison of contact frequencies between the two ORCA biological replicates at a contact threshold of 150 nm. Pearson’s R^2^ shows the correlation between the contact frequencies of both replicates. **(e) (Left)** Upset plot summarizing the multi-way interaction frequency between regulatory elements such as CTCF binding sites found to be most interacting based on wildtype DP T cell HiC. Schematic shows the interaction network highlighting the most interacting 5-way node network. Red bar denotes the number of alleles which do not interact with any of the other nodes. **(Right)** Allele frequency measured as a function of the number of interacting nodes within the interaction network in wildtype DP T cells. ORCA data from the *Sox2* locus^37^ used as a control to quantify the interaction frequency of 5 distance-matched nodes. **(f)** Distribution of *Gapdh* mRNA count across different fields of view (FOVs) imaged. Significance computed using two-sided Wilcoxon rank sum test. **(g)** Representative chromatin traces reconstructed using ORCA showing the three-way interaction hub between *Fli1*, *Ets1* and *Gm27162* **(top)** along with an example showing *Fli1-Ets1* promoter-promoter interaction **(bottom)** within the same population of wildtype DP T cells. Distances of each interacting pair are shown beside the reconstructed chromatin traces. Color bar shows the pseudo-color assigned to the 30 kbp readout sequences used to sequentially image the *Ets1-Fli1* locus.

**Figure S3.**
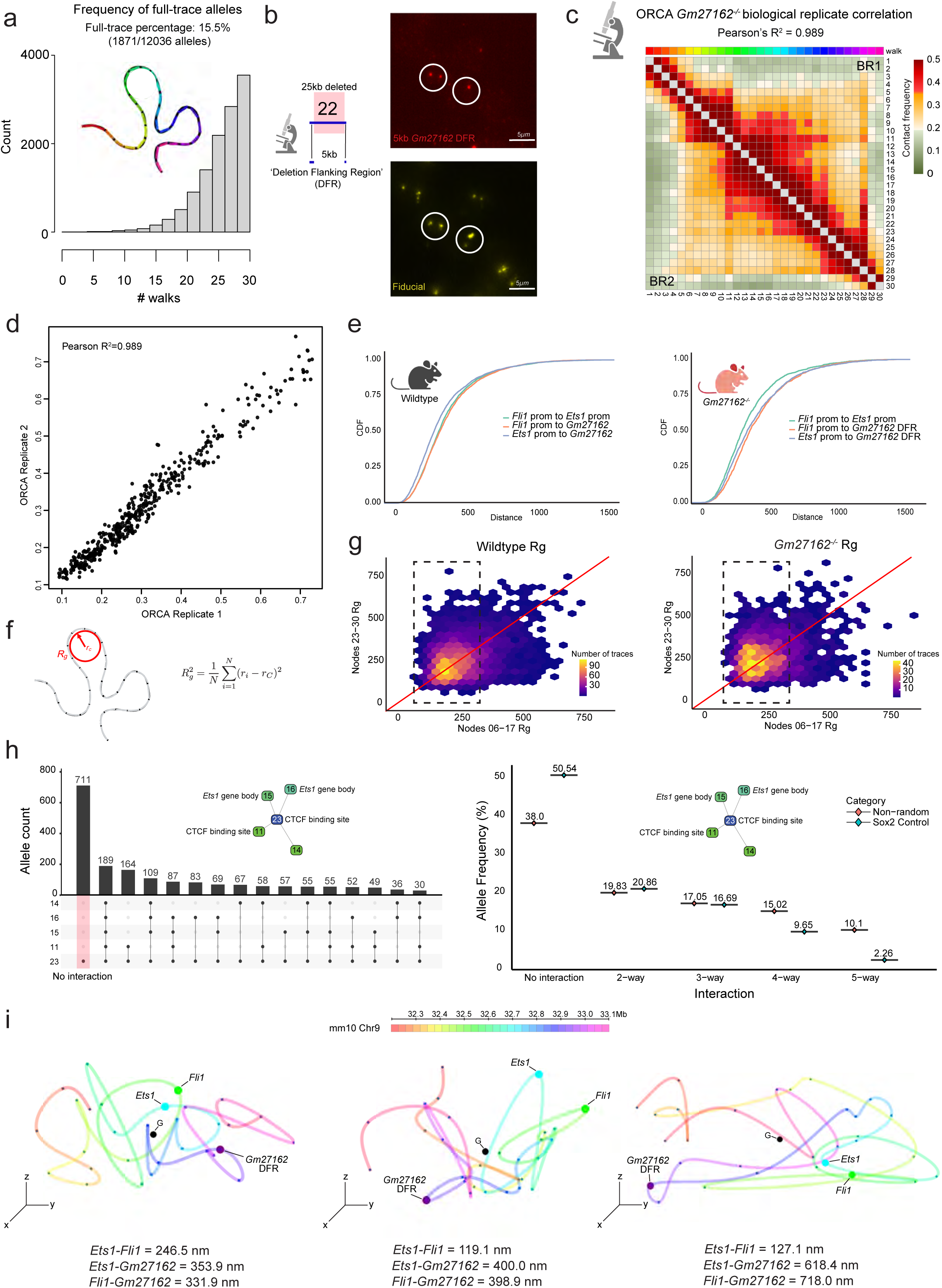
Deletion of the *Ets1* super-enhancer *Gm27162* increases promoter-promoter interactions between *Ets1* and *Fli1*. (a) Bar plot displays the proportion of *Gm27162*^-/-^ DP T cell chromatin traces as a function of the number of readouts successfully hybridized. Schematic of chromatin trace with all 30 successful hybridizations shown along with full-trace percentage. (b) Representative images showing successful hybridization of ORCA readout probe **(top)** on the 5 kbp region flanking *Gm27162* in *Gm27162*^-/-^ DP T cells co-localizing with fiducial reference signal **(bottom)**. Schematic on the left shows that the 25 kbp deletion falls within a single 30 kbp ORCA bin, with the remaining 5 kbp fragment defined as the ‘Deletion Flanking Region’ (DFR). (c) Correlation between two ORCA *Gm27162*^-/-^ biological replicates displayed as Pearson’s R^2^ value. Traces with all 30 successful hybridizations were used to generate the distance matrix, with distances corresponding to readout 22 coming from the imaging and detection of the 5 kbp *Gm27162* DFR. (d) Comparison of contact frequencies between the two *Gm27162*^-/-^ DP T cell ORCA biological replicates at a contact threshold of 150 nm. Pearson’s R^2^ shows the correlation between the contact frequencies of both replicates. (e) Cumulative distance distribution between the promoters of *Fli1* to the promoter of *Ets1* (green, wildtype mean = 369.7 nm, *Gm27162*^-/-^ mean = 368.2 nm), promoter of *Fli1* to *Gm27162* (orange, wildtype mean = 377.1 nm, *Gm27162*^-/-^ mean = 425.6 nm) and promoter of *Ets1* to *Gm27162* (blue, wildtype mean = 342.1 nm, *Gm27162*^-/-^ mean = 414.6 nm) in wildtype DP T cells **(left)** and *Gm27162*^-/-^ DP T cells (**right).** (f) Schematic of chromatin trace used to show the computation of the radius of gyration (Rg, red circle) about the geometric center of each ORCA chromatin trace. Equation used to compute Rg is shown where ri represents the (x,y,z) position of readout ‘i’ and rc represents the geometric center for a given chromatin trace (g) Scatterplot comparing the compaction of the acetylated chromatin between *Ets1-Fli1* (x-axis) and flanking methylated (y-axis) compartment in wildtype **(left)** and *Gm27162*^-/-^ **(right)** DP T cells measured by radius of gyration (Rg). Dashed rectangular box shows the enrichment of chromatin traces with more compact *Ets1-Fli1* domains in *Gm27162*^-/-^ relative to wildtype DP T cells. (h) **(Left)** Upset plot summarizing the multi-way interaction frequency between regulatory elements such as CTCF binding sites found to be most interacting based on wildtype DP T cell HiC. Schematic shows the interaction network highlighting the most interacting 5-way node network. Orange bar denotes the number of alleles which do not interact with any of the other nodes. **(Right)** Allele frequency measured as a function of the number of interacting nodes within the interaction network in *Gm27162*^-/-^ DP T cells. ORCA data from the *Sox2* locus^37^ used as a control to quantify the interaction frequency of 5 distance-matched nodes. (i) Representative ORCA chromatin traces showing decreased proximity between *Fli1* and *Ets1* along with the ejection of the *Gm27162* DFR in *Gm27162*^-/-^ DP T cells. Distances of each interacting pair are shown below the reconstructed chromatin traces. Color bar shows the pseudo-color assigned to the 30 kbp readout sequences used to sequentially image the *Ets1-Fli1* locus.

**Figure S4.**
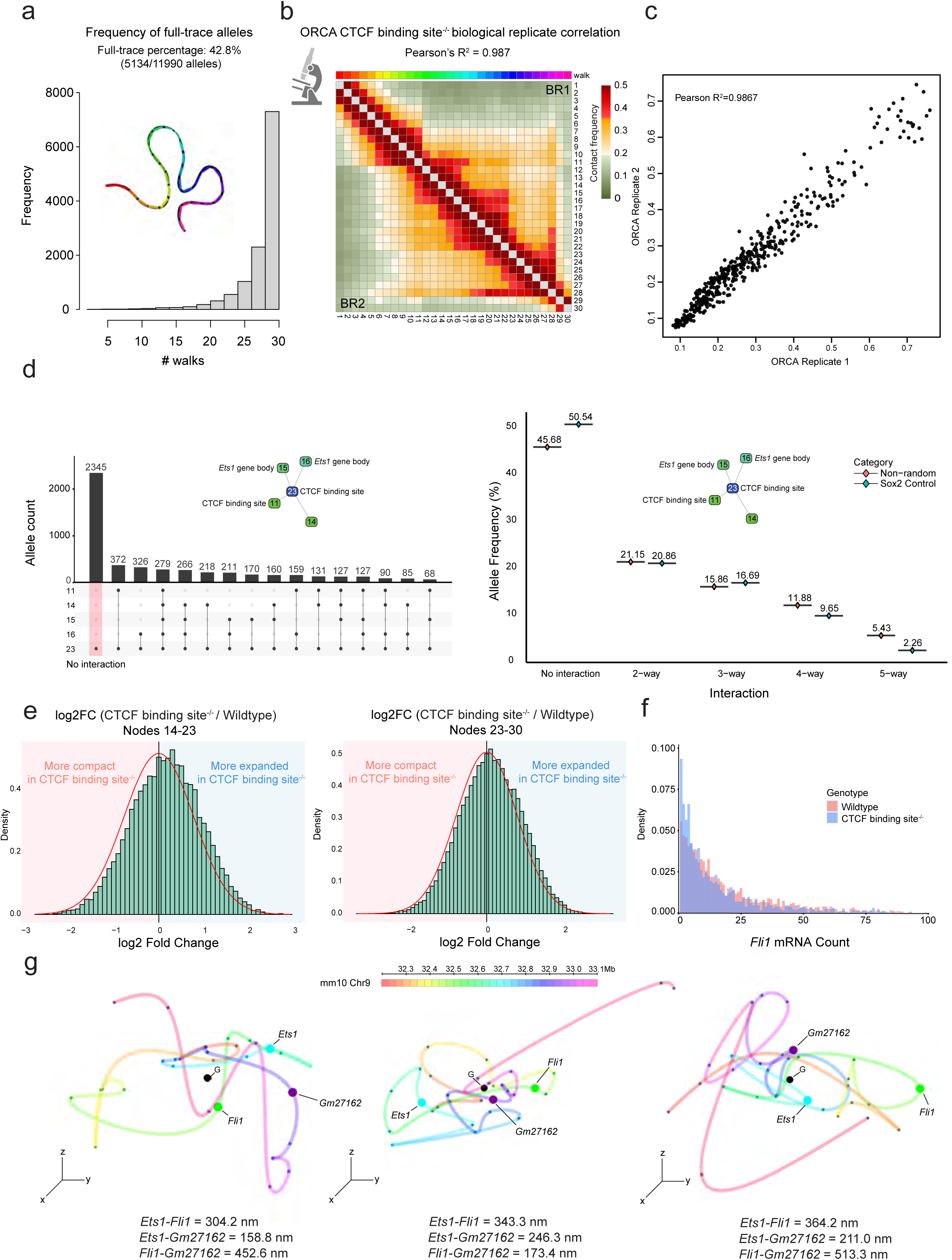
Deletion of the CTCF binding site at the interface between H3K27ac-H3K27me3 transition causes domain-wide decompaction at the *Ets1-Fli1* locus. (a) Bar plot displays the proportion of CTCF binding site^-/-^ DP T cell chromatin traces as a function of the number of readouts successfully hybridized. Schematic of chromatin trace with all 30 successful hybridizations shown along with full-trace percentage. (b) Correlation between two CTCF binding site^-/-^ ORCA biological replicate experiments displayed as Pearson’s R^2^ value. (c) Comparison of contact frequencies between the two CTCF binding site^-/-^ DP T cell ORCA biological replicates at a contact threshold of 150 nm. Pearson’s R shows the correlation between the contact frequencies of both replicates. (d) **(Left)** Upset plot summarizing the multi-way interaction frequency between regulatory elements such as CTCF binding sites found to be most interacting based on wildtype DP T cell HiC. Schematic shows the interaction network highlighting the most interacting 5-way node network. Orange bar denotes the number of alleles which do not interact with any of the other nodes. **(Right)** Allele frequency measured as a function of the number of interacting nodes within the interaction network in CTCF binding site^-/-^ DP T cells. ORCA data from the *Sox2* locus^37^ used as a control to quantify the interaction frequency of 5 distance-matched nodes. (e) Histogram of log2(fold change) difference in the compaction of the acetylated domain upstream of the *Ets1* gene (i.e., between readouts 14-23) **(left)** and methylated domain downstream of the CTCF binding site deletion (i.e., between readouts 23-30) **(right)** between CTCF binding site^-/-^ and wildtype DP T cells. The y-axis represents the frequency of observing a particular log fold change which was computed by bootstrapping 1000 alleles per genotype for 1000 iterations and calculating the local Rg ratio at each iteration to account for differences in total allele count between CTCF binding site^-/-^ DP T cells and wildtype DP T cells. The red area corresponds to more compact chromatin structure whereas the blue area corresponds to a more expanded chromatin structure. In both histograms, the red curve represents the shape of the distribution if it was normally distributed. (f) Density histogram comparing the levels of *Fli1* mRNA counts measured by single molecule RNA FISH in CTCF binding site^-/-^ (blue bars) and wildtype DP T cells (red bars). (g) Representative ORCA chromatin traces showing preserved interactions between *Ets1-Gm27162* and an increase in distances between *Fli1* and *Ets1*. Distances of each interacting pair are shown below the reconstructed chromatin traces. Color bar shows the pseudo-color assigned to the 30 kbp readout sequences used to sequentially image the *Ets1-Fli1* locus.

**Figure S5.**
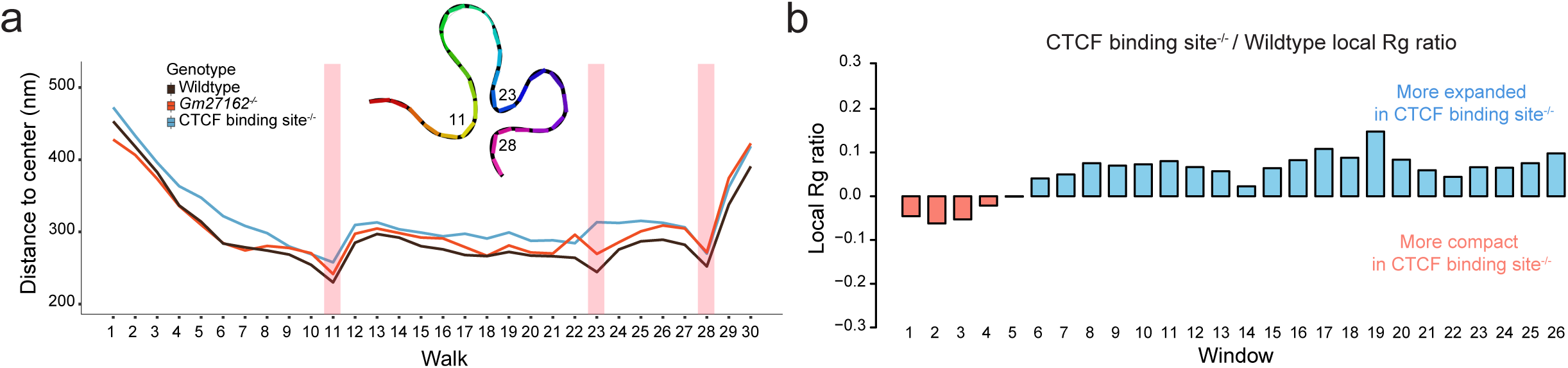
Perturbing H3K27ac-H3K27me3 balance increases domain-wide disorder in chromatin folding. **(a)** Mean position of each readout plotted relative to the geometric center of each trace for wildtype (black), *Gm27162*^-/-^ (red) and CTCF binding site^-/-^ (blue) DP T cells. Schematic shows the chromatin organization mediated by the radial positioning of CTCF binding sites within the *Ets1-Fli1* multi-enhancer hub. **(b)** Local radius of gyration (Rg) computed across the *Ets1-Fli1* multi-enhancer hub using 5 consecutive readout window shifted along the entire locus. Barchart shows the ratio of local Rg between CTCF binding site^-/-^ and wildtype DP T cells. Bars shaded red represent more compact chromatin domains whereas bars shaded blue represent less compact (or more expanded) chromatin domains

